# Role of RSPO3 in Estrogen-mediated Sex Differences in Body Fat Distribution: A Single-cell Data-driven Study

**DOI:** 10.1101/2025.01.03.631121

**Authors:** Qian Xu, Yongjian Yang, Shreyan Gupta, Bo-Jia Chen, Guanxun Li, Mathew Chamberlain, James J Cai

## Abstract

Obesity is a global health concern affecting more than one-third of the adult population worldwide. Epidemiological studies indicate that body fat distribution, more than overall adiposity, is a key risk factor for obesity-related diseases. Genome-wide association studies revealed that RSPO3 is associated with body fat distribution. Additionally, marked sex differences in body fat distribution are largely influenced by estrogen, suggesting that exploring the relationship between estrogen and RSPO3 expression may illuminate the mechanisms underlying the sexual dimorphism of fat distribution. We revisited a comprehensive set of gene expression data from humans and mice to explore these mechanisms. We assessed RSPO3 expression in male and female adipose tissues and utilized single-cell RNA sequencing (scRNA-seq) and spatial transcriptomics to examine RSPO3’s function within different tissues. Our findings confirmed significantly higher RSPO3 expression in male adipose tissue compared to female. Furthermore, our data for the first time suggest that estrogen plays a crucial role in fat distribution, as blocking estrogen signaling upregulates RSPO3 expression. In mouse liver, we observed a correlation between sex-specific ADK overexpression and RSPO3 expression, resulting in increased obesity and hepatocyte expansion via WNT/β-catenin signaling in male mice. This study sheds light on how estrogen influences sexual dimorphism in fat distribution and underscores RSPO3’s vital role in this process. These insights open new avenues for targeted therapeutic interventions addressing obesity and metabolic diseases with a focus on sex-specific factors.

## 1. Introduction

Obesity is an epidemic disease that threatens the ability to inundate healthcare resources by increasing the incidence of diabetes, heart disease, hypertension, and cancer [1]. While obesity is traditionally measured by body mass index (BMI), this metric does not account for fat distribution in various body compartments. An alternative measure, the waist-to-hip ratio, offers insights into this body fat distribution. Central obesity, characterized by an accumulation of visceral adipose tissue (VAT), has been associated with a heightened risk of metabolic disturbances, encompassing insulin resistance and Metabolic dysfunction-associated steatotic liver disease (MASLD), and malignancies such as hepatocellular carcinoma (HCC) [2]. In contrast, increased peripheral subcutaneous adipose tissue (SAT) accumulation may confer protection against these disorders [3].

Important sexual dimorphism in body fat distribution has been the focus of various studies [2, 4, 5]. Typically, women accumulate more SAT, resulting in a “pear shape”, while men tend to accumulate more VAT, leading to an “apple shape”. This sexual dimorphism diminishes post-menopause [6] as female hormones decrease, leading to a rise in abdominal adiposity, similar to that observed in men [7]. Studies characterizing sexual dimorphism in body fat distribution have unsurprisingly pointed to sex hormones, particularly estrogens [8-10]. Estrogens are believed to play a protective role in obesity-related diseases, such as diabetes and HCC, which potentially explains the correlation between body fat distribution and the risk of obesity-related diseases [11]. Although the protective role of estrogen is supported by these lines of evidence, the underlying mechanisms remain unclear.

Genome-wide association studies (GWASs) have identified approximately 20 loci that are linked to body fat distribution [12-14]. Among these, the *RSPO3* gene, known for its oncogenic potential [15], has garnered significant attention. In addition, *RSPO3* is implicated in determining peripheral adipose tissue storage capacity and influences body fat distribution by regulating adipose cell biology [16]. Previous studies have also shown higher RSPO3 expression in male adipose tissue [16]. Specifically, *RSPO3* promotes upper-body fat distribution by stimulating *abdominal* adipose progenitor proliferation while inhibiting *gluteal* adipose progenitor differentiation [16]. It is known that *RSPO3’s* actions are associated with changes in the WNT signaling pathway. The regulatory mechanisms governing *RSPO3* expression, its role in cell-cell communication, and its downstream effects on target tissues remain largely unexplored.

A past study [17] in adenosine kinase (*ADK*) transgenic mice demonstrated that overexpressing *ADK* can induce obesity. *ADK* encodes the enzyme that phosphorylates adenosine to generate adenosine monophosphate. Recent studies suggested a role for *ADK* in regulating fat metabolism and systemic insulin sensitivity. Of note, a study by Boison *et al.* [18] revealed increased liver fat content in neonatal mice on global *ADK* disruption, and a study by Xu *et al.* [19] showed decreased severity of high-fat diet-induced obesity and hepatic steatosis in mice as a result of the endothelial *ADK* disruption. Li *et al.* [17] also observed that hepatocyte ADK functions to promote excessive fat deposition and liver inflammation. They also noted a sex-based discrepancy in the effects of *ADK* overexpression: while it induced obesity in males, females exhibited a more subdued response. Both male and female *ADK* overexpressing mice (fed a standard chow diet) exhibited greater body weight, greater adiposity, more severe hepatic steatosis, and greater systemic insulin resistance than control mice. However, female mice experienced a slower onset of these symptoms—the onset of weight gain and fat accumulation in female mice was more gradual [17]. Furthermore, *ADK* overexpression female mice exhibited less pronounced insulin resistance and maintained steadier blood glucose levels than did their male counterparts [17]. This relative protection of female mice from the obesogenic effects of ADK prompted us to investigate if the estrogen-*RSPO3* axis plays a role in this sex difference in the ADK-induced mouse obesity model.

This study presents a comprehensive single-cell level analysis of the molecular mechanisms underlying estrogen-and *RSPO3*-mediated sex differences in body fat distribution. Our findings suggest a mechanistic link between estrogen and *RSPO3* signaling, thus, offering novel insights into the biological basis of sexual dimorphism of adipose tissue patterning.

## 2. Methods

### 2.1. Expression data processing

In this study we used a human microarray dataset from Icelandic Family Adipose (IFA) cohort (GSE7965) and a bulk RNA-seq dataset of adipose tissues from the GTEx database to conduct tissue-level analyses. For single-cell level analyses, we used two single-cell RNA-seq (scRNA-seq) datasets from mouse liver and breast tissues (GSE191219), and one spatial transcriptomics dataset from human colorectal cancer tissue.

The microarray data from the Icelandic Family Adipose (IFA) cohort (*n* = 701), including 297 males and 404 females, was analyzed using the R software environment (v4.2.2) with the Limma package (v3.54.2). The sex of samples was determined based on a threshold of -0.2 in XIST gene expression. Subsequently, gene expression data was normalized using the “normalizeQuantiles” function. A design matrix was created to model the impact of sex on gene expression, with ‘Male’ and ‘Female’ as two levels. Differential gene expression between the sexes was estimated using “lmFit” and “contrasts.fit” functions. Empirical Bayes smoothing was applied using the “eBayes” function. The list of top differentially expressed genes was generated using the “topTable” function, with *p*-values adjusted using the Benjamini-Hochberg method to control the false discovery rate. This output was used to identify the most significantly differentially expressed genes between males and females in the cohort. The scRNA-seq data was processed in the R software environment (v4.2.2) using the Seurat package (v4.3.0). Expression matrices from VCD_Vehicle, VCD_E2, Intact, and VCD_E2_ICI were individually loaded into R using the “Read10X” function and subsequently utilized to construct Seurat objects. The “NormalizeData” function was utilized to adjust the number of unique molecular identifiers (UMIs) in each cell, followed by data scaling using the “ScaleData” function. Principal Component Analysis (PCA) was carried out on the scaled data using the RunPCA function to reduce data dimensionality. For visualization of high-dimensional cellular data, Uniform Manifold Approximation and Projection (UMAP) was implemented via the “RunUMAP” function. Cells were clustered using the “FindNeighbors” and “FindClusters” functions in Seurat. Violin plots and UMAP plots were used for visualization of gene expressions and clusters, respectively. The expression of gene *RSPO3* was particularly noted and compared across groups using a box plot constructed with “ggboxplot” and statistical comparison added using “stat_compare_means”. All parameters were set to default.

### 2.2. Differential expression analysis against synthetic null data

Differential expression (DE) analysis of the scRNA-seq data was performed using the ClusterDE package (v0.99.0). We first performed a DE test on the original data for the two clusters we were interested in to identify DE genes. The test used the Wilcoxon rank-sum test from the ‘FindMarkers’ function in the Seurat package. We then generated synthetic null data based on the original data which contains the two clusters we were interested in, such as the “WT” and the “OE” group of cluster 2 hepatocyte, using the “constructNull” function. Then, we performed a standard Seurat analysis using the synthetic null data, including normalization and clustering. We then performed a DE test on the synthetic null data using the Wilcoxon rank-sum test. Then we extracted the *p*-values of the DE genes from synthetic null data and compared them with the *p*-values of the DE genes from the original data using the “ClusterDE” function. This function used the FDR-control method, Clipper [20] to calculate a contrast score based on the two DE scores for each gene.

### 2.3. Spatial transcriptomics data analysis

The spatial transcriptomics data analysis was performed using the Seurat package. The 10x Genomics Visium spatial transcriptomics data was loaded using the “Load10X_Spatial” function. The data were normalized, and variance-stabilizing transformation was performed using the “SCTransform” function. PCA was conducted using the “RunPCA” function to reduce the dimensionality of the data. To identify the clusters of cells with similar transcriptomic profiles, we used the “FindNeighbors” and “FindClusters” functions. The UMAP plot was produced using the “RunUMAP” function to visualize the high-dimensional data in a low-dimensional space. Specific gene expression on the UMAP plot and spatial plots was visualized using “FeaturePlot” and “SpatialFeaturePlot” functions respectively. For each cluster, differentially expressed genes were identified using the “FindAllMarkers” function. The Nebulosa package (v1.8.0) was employed to generate spatial density plots for specific genes of interest.

## 3. Results

### 3.1. Higher RSPO3 expression in male adipose tissue than in female adipose tissue

To explore potential differences in *RSPO3* expression related to sex, we conducted an analysis using RNA-seq data from the GTEx database [21]. This dataset is composed of subcutaneous adipose tissue samples from 663 donors and visceral adipose tissue samples from 541 donors. Our investigation revealed consistent variations in *RSPO3* expression in the two adipose tissue subtypes between males and females. In both subcutaneous and visceral adipose tissues, females exhibited lower *RSPO3* expression levels than males did (**Fig. 1A**). To substantiate these findings, we utilized microarray data from a prior study involving 701 samples of human subcutaneous adipose tissue [22]. This analysis further confirmed our initial observations, demonstrating significantly greater *RSPO3* expression in males than in females (log2 fold change = 0.16, *p* = 1.48e-27; **Fig. 1B**).

**Figure 1.**
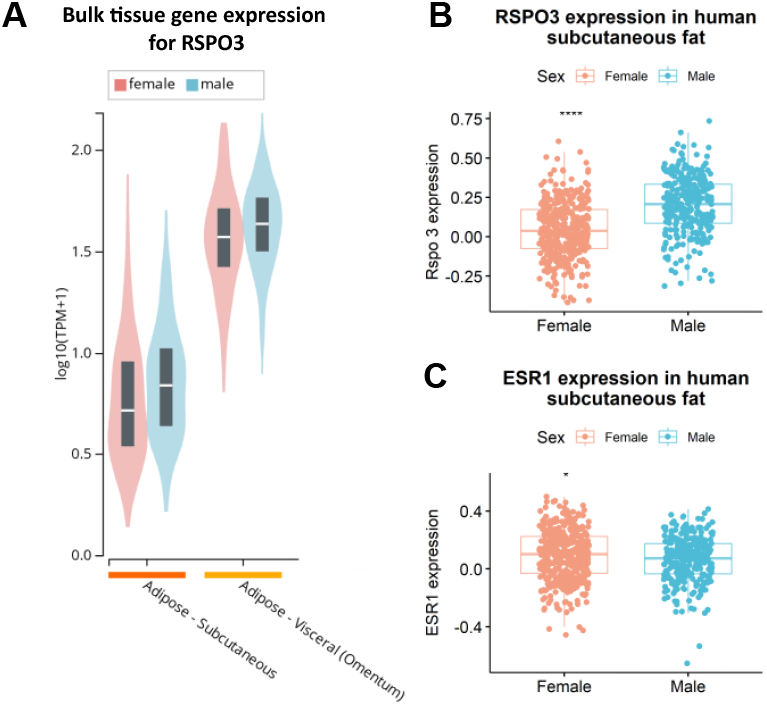
Differential *RSPO3* and *ESR1* expression in human adipose tissue between males and females. (**A**) *RSPO3* expression level in human subcutaneous and visceral adipose tissue. Data source: GTEx [21]. (**B**) *RSPO3* expression in female and male subcutaneous adipose tissue. Data source: GEO GSE7965. (**C**) *ESR1* expression in female and male subcutaneous adipose tissue. Data source: GEO GSE7965.

Thus, our analyses highlighted a disparity in *RSPO3* expression between males and females. Given the association of *RSPO3* with body fat distribution and the waist-to-hip ratio, the elevated *RSPO3* levels in male adipose tissue may offer insights into the sexual dimorphism observed in body shape. Furthermore, we assessed the expression levels of *ESR1*, an estrogen receptor, in the same microarray dataset. ESR1 exhibits slightly but significantly higher expression levels in females (log2 fold change = 0.024, *p* = 0.045; **Fig. 1C**). Since females typically possess higher estrogen levels than males, these findings suggest that estrogen may play a role in modulating *RSPO3* gene expression, potentially elucidating the observed variations in body fat distribution between the sexes.

### 3.2. RSPO3 expression is suppressed by estrogen

Building on our observations of differences in *RSPO3* expression in male and female human adipose tissues, we further explored the influence of estrogen on *RSPO3* expression. To this end, we utilized a scRNA-seq dataset from a female mouse mammary gland model treated with 4-vinylcyclohexene diepoxide (VCD), which simulates menopausal conditions, without affecting other tissues [23, 24]. The mammary gland model was used due to its high *RSPO3* expression, estrogen sensitivity, relevance to body fat distribution, and accessibility. This model allowed for the investigation of *RSPO3* regulation in a physiologically relevant context, leading to insights into the molecular mechanisms underlying sexual dimorphism in fat distribution.

The mice were subjected to three different treatments: vehicle, estrogen (E2), or a combination of E2 and ICI, which is a steroidal estrogen antagonist that was designed to be devoid of estrogen agonist activity in preclinical models [25]. An untreated group served as the control (Intact group). Our primary objective was to ascertain the impact of the estrogen-estrogen receptor signaling pathway on *RSPO3* expression by comparing the expression levels of *RSPO3* across the four different treatment conditions: Vehicle, E2, E2 combined with ICI, and Intact.

To identify the cell types in this dataset and determine the source of *RSPO3* expression, we first visualized all cells with UMAP **(Fig. S1A**). The UMAP plot showed an intermingling of cells from different treatment groups, forming 23 distinct clusters (**Fig. S1B**). We annotated each cluster based on marker gene expression, referencing previous studies [23, 26, 27] and methods [28, 29]. This analysis revealed fifteen cell types, including six fibroblast subgroups and various immune cell types (**Fig. 2A**), with the marker genes detailed in **Fig. 2B**.

**Figure 2.**
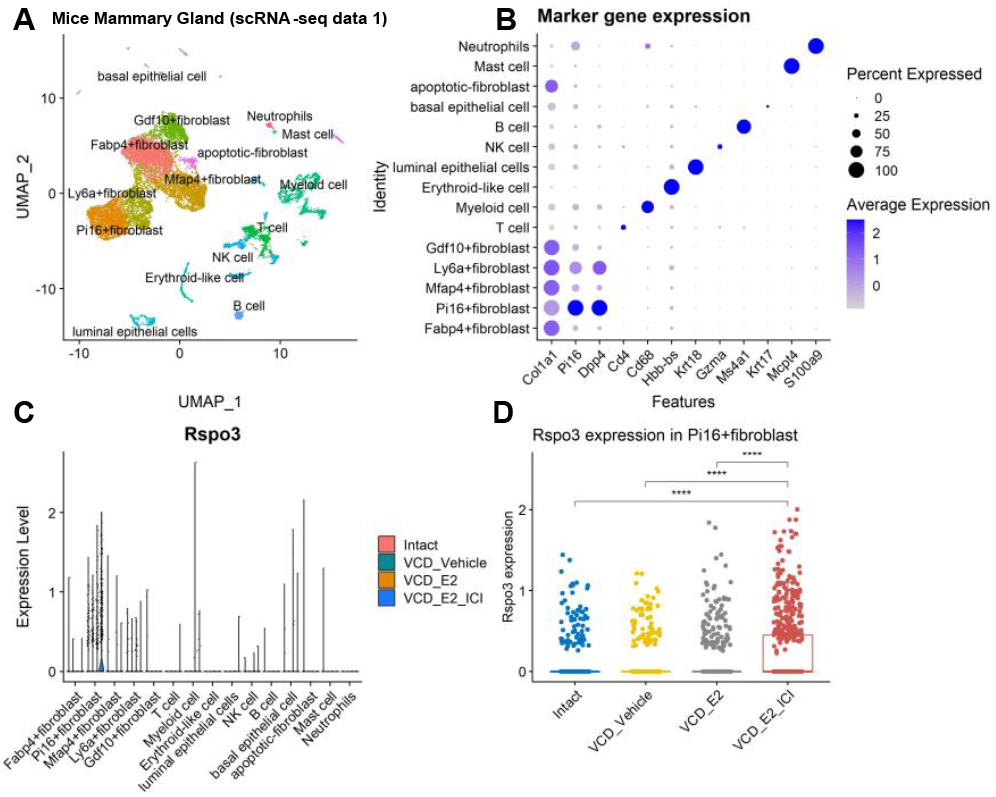
Estrogen-mediated regulation of *RSPO3* expression in mouse mammary glands. (**A**) UMAP visualization of diverse cell populations within the mouse mammary gland. (**B**) Marker genes for the six identified fibroblast clusters, as well as the other cell types. (**C**) *RSPO3* expression in different cell types. The six fibroblast clusters are underlined; Pi16+ fibroblasts had the highest *RSPO3* expression among all the fibroblasts. (**D**) Comparison of *RSPO3* expression across four treatment groups: A comparative analysis showing *RSPO3* expression levels in the four treatment groups: Intact, VCD_Vehicle, VCD_E2, and VCD_E2_ICI.

A thorough examination of mouse mammary gland scRNA-seq data revealed that the primary source of *RSPO3* was the Pi16+ fibroblast cluster (**Fig. 2C**). This cluster, which is identified as *Pi16*+ *Dpp4*+ fibroblasts, resembled adventitial stromal cells typically found in vascular settings [30]. These findings align with the existing knowledge of *RSPO3* as an endothelial-specific angiocrine factor [31].

We next focused on the potential impact of estrogen on *RSPO3* expression patterns. We observed that fibroblasts from the VCD_E2_ICI group (*n* = 705), where the estrogen effect was neutralized, exhibited significantly higher *RSPO3* expression compared to those from the Intact group (*n* = 721, *p* = 1.2e-05) and the VCD_E2 group (*n* = 499, *p* = 6.7e-09) (**Fig. 2D**). This indicated that the elimination of estrogen-estrogen receptor signaling led to increased *RSPO3* expression in the mammary glands of these female mice.

From these analyses, we draw three key conclusions: (1) *RSPO3* expression is highly specific to the cell type and occurs predominantly in a distinct subset of fibroblasts; (2) the estrogen-estrogen receptor signaling pathway exerts an inhibitory influence on *RSPO3* expression; and (3) the estrogen receptor plays a crucial role in the estrogen-mediated downregulation of *RSPO3* expression. These findings contribute to our understanding of the molecular relationship between estrogen signaling and *RSPO3*, as well as their potential role in influencing sexual dimorphism in body fat distribution.

**Figure S1.**
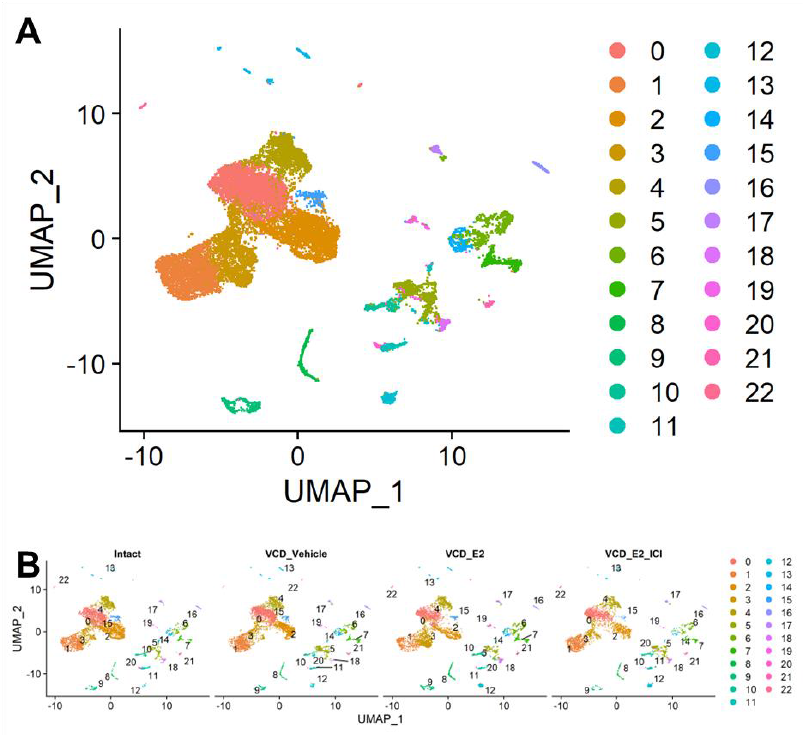
UMAP visualization of the clusters from murine mammary glands. (**A**) Twenty-three clusters in the mouse mammary gland scRNA-seq dataset. (**B**) The same UMAP plot split into 4 different treatment groups. The cells from the different treatment groups were mixed well with each other.

### 3.3. RSPO3 is expressed differently in different cell types and functions via a paracrine mechanism

Our initial observations revealed that *RSPO3* expression in the mouse mammary gland is highly cell-type specific and occurs primarily in a small subset of fibroblasts. This was evidenced by our analysis of the *mouse mammary gland scRNA-seq dataset* (**Fig. 2C**). This difference in expression patterns led us to hypothesize that *RSPO3* may function through a paracrine mechanism.

To test this hypothesis, we examined two additional datasets: (1) *10x Visium data*—a spatial transcriptomics dataset procured from human colorectal cancer, and (2) *liver NPC scRNA-seq data*—a dataset from mouse liver nonparenchymal cells (NPCs) [17]. We aimed to validate the location of *RSPO3* expression in tissues and identify any prospective paracrine interactions between cell types.

The *10x Visium data* from human colorectal cancer patients included 9,080 spots, and the cell types were delineated based on specific marker gene expression (**Fig. 3A, C**). We identified six cell types, namely, B/plasma cells (*IGHD*+), epithelial cells (*CA9*+, *PIGR*+, *FCGBP*+, and *MUC5B*+), stromal cells (*NQO1*+), myeloid cells (*CXCL8*+), and T and NK cells, as well as innate lymphoid cells (*TNKILC*, marked by *SLC7A5*+). The marker genes used for cell type identification were cross-verified by referring to the markers in the Human Colon Cancer Atlas [32].

**Figure 3.**
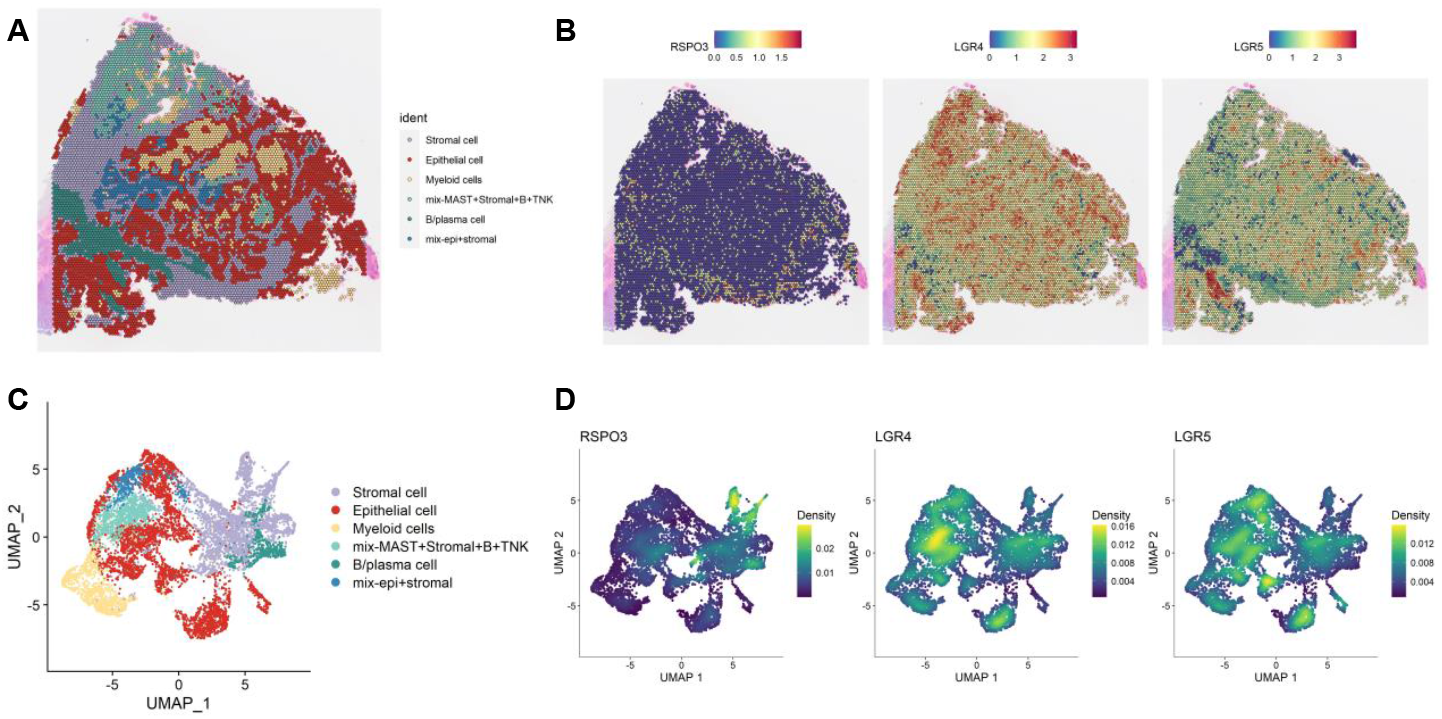
Spatial transcriptomics reveals the paracrine mode of action of *RSPO3* in human colorectal cancer. (**A**) Visualization of the primary cell type under each spot. The 10x Visium spatial transcriptomics data-derived cell type labels are superimposed on the image, highlighting six primary cell types: Stromal cells, epithelial cells, myeloid cells, a mixed cluster of epithelial and stromal cells, B/plasma cells, and a mixed cluster of stromal and immune cells (MAST+stromal+B+TNK). (**B**) Spatial expression of *RSPO3* and two selected receptor genes, *LGR4/5. RSPO3* is the ligand that is expressed mainly in stromal cells (left), while the receptors LGR4 (middle) and *LGR5* (right) are expressed predominantly in a mixed population of immune and stromal cells and epithelial cells, respectively. (**C**) UMAP visualization. A UMAP representation of the 10x Visium data delineating the six distinct cell type components. (**D**) UMAP plot of *RSPO3* and *LGR4/5* expression. *RSPO3* was expressed mainly in stromal cells, *LGR4* was expressed mainly in epithelial cells (middle), and *LGR5* was expressed mainly in epithelial cells (right).

Our analysis of the *10x Visium data* paralleled our findings in the mouse mammary gland: *RSPO3* expression is highly specific to certain cell types, and the spatial distribution of *RSPO3*-expressing cells is distinct from that of cells expressing its receptor LGR4/5 (**Fig. 3B**). Specifically, *RSPO3* was expressed mainly by stromal cells (**Fig. 3B**, left panel; **Fig. 3D**, left panel), while the receptor LGR4 was expressed predominantly in a mixture of stromal and immune cells (labeled MAST+stromal+B+TNK cell types), and *LGR5* was expressed in epithelial cells (**Fig. 3B,** middle and right panels; **Fig. 3D,** middle and right panels). These findings further support our earlier conclusion regarding the cell type-specific expression of *RSPO3* and its paracrine mode of action.

We then evaluated the *liver NPC scRNA-seq data* to confirm the paracrine signaling mechanism of *RSPO3*. This dataset included nine cell types in addition to hepatocytes (*LGN*+), each characterized by distinct markers [17] (**Fig. 4A**), including T cells (*TRBC1*+), erythroid-like cells (*HBB-BS*+), neutrophils (*S100A8*+), hepatic stellate cells (HSCs, *DCN*+), endothelial cells (*ENG*+), B cells (*CD79A*+), monocytes (*CCR2*+), macrophages (*MPEG1*), and cholangiocytes (*SOX9*+).

**Figure 4.**
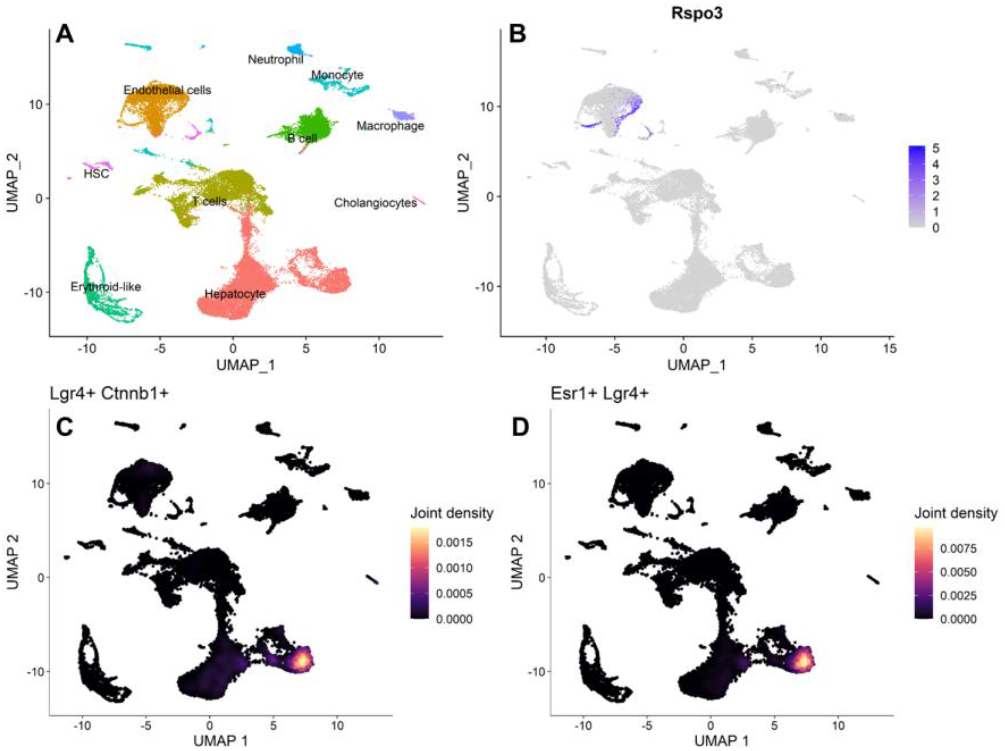
Increased *Rspo3* expression in liver endothelial cells due to ADK overexpression. (**A**) Depiction of the various cell types present in mouse liver NPCs. (**B**) Feature plot highlighting that *Rspo3* expression originates predominantly from endothelial cells. (**C**) Joint density plot illustrating the pronounced difference in the expression of the receptor gene, *Lgr4*, and the β-catenin gene, *Ctnnb1*, in the WNT signaling pathway in hepatocytes. (**D**) Joint density plot indicating co-expression of *Esr1* and *Lgr4* in hepatocytes.

Our analysis revealed that *RSPO3* expression in liver NPCs was mainly localized to endothelial cells (**Fig. 4B**) and not to hepatocytes or immune cells. Interestingly, the *RSPO3* receptor *LGR4* was exclusively found in hepatocytes (**Fig. 4C, D**). These results align with our initial hypothesis that *RSPO3*-*LGR4* signaling is highly cell-type specific, further confirming the paracrine mode of action of *RSPO3*. In the context of the liver, *RSPO3* is secreted by endothelial cells and impacts adjacent *LGR4*+ hepatocytes.

### 3.4. ADK overexpression increases RSPO3 expression in mice liver

We focused on the *liver NPC scRNA-seq data* to study the effect of *RSPO3* signaling. This dataset included two groups of samples: the ADK overexpression (ADK-OE) group and the control (WT) group. The ADK-OE group consisted of male mice with hepatocyte-specific ADK-OE mutation, while the control group included male mice treated with an empty AAV vector. By comparing these two groups, we observed an increased *Rspo3* expression level in the ADK-OE group compared to the control group (**Fig. 5A**). The endothelial cells in the ADK-OE group showed significantly higher *Rspo3* expression compared with the endothelial cells in the control group (*t*-test, *p* < 0.0001) and HSCs (*t*-test, *p* < 0.0001) in the liver. Although the dataset lacked data from female mice, our previous results on the mouse mammary gland data suggest that estrogen may suppress *Rspo3* expression. Considering this, along with the fact that female mice are relatively protected from the obesogenic effects of *ADK*, we hypothesize a protective role for estrogen in the liver, mitigating fat accumulation and maintaining insulin sensitivity by inhibiting *Rspo3* expression. In contrast, male mice, which lack sufficient estrogen, display a more pronounced response to ADK-OE-induced *Rspo3* expression, leading to more severe obesity and systemic insulin resistance.

**Figure 5.**
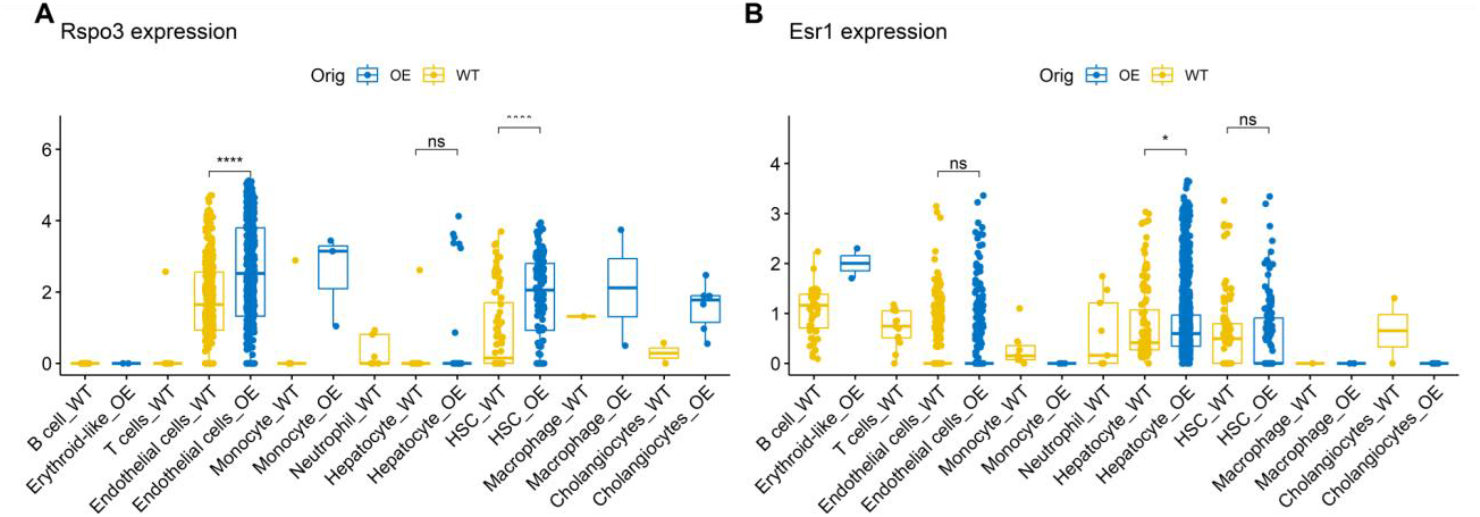
Increased *Rspo3* and Esr1 expression in endothelial cells and hepatocytes due to ADK overexpression. (**A**) Box plot showing *Rspo3* expression across various cell types in WT and ADK-OE samples. Notably, endothelial cells and HSCs in the ADK-OE samples show a significant increase in *Rspo3* expression (*t*-tests, both *p* < 0.0001****). (**B**) Box plot showing Esr1 expressions across various cell types. Hepatocytes in the ADK-OE sample show a significant increase in Esr1 expression (*t*-test, *p* < 0.01*).

We also noted a heightened expression of the estrogen receptor gene *Esr1* in hepatocytes from the ADK-overexpression group (**Fig. 5B,** *t*-test, *p* < 0.0001). It has previously been established that *Rspo1*, another R-spondin protein, can enhance *Esr1* expression, thereby modulating the activity of steroid hormone signaling in the mouse mammary gland [33]. Our findings suggest that *Rspo3* may likewise influence *ESR1* expression in hepatocytes, potentially acting as a modulator of estrogen signaling.

### 3.5. RSPO3 drives hepatocyte expansion, influences metabolic processes, and activates oncogenic pathways

Enhanced expression of *RSPO3* has been discovered in colon cancer patients, highlighting its potential role in colorectal cancer development. Although *RSPO3*’s overactivation and its oncogenic effects are well-documented, the majority of RSPO-related cancer research focuses on the intestinal tract [34-37]. Yet, the specific oncogenic role of *RSPO3* in the liver remains less understood. To address this gap, we examined *RSPO3* signaling effects in liver cells. Given *RSPO3*’s established role in promoting *LGR5*+ intestinal stem cell expansion and subsequent intestinal tumorigenesis, we postulated that *RSPO3* might exert a comparable influence on liver cells, possibly through similar mechanisms involving the WNT signaling pathway.

In our detailed analysis of the *liver NPC scRNA-seq data*, we focused specifically on hepatocytes due to the predominant expression of *LGR4*, a key *RSPO3* receptor, in these cells. By comparing the ADK-OE group and the control group, we made two key findings. (1) In the ADK-OE group, the *LGR4*+ /β-catenin+ (*CTNNB1*) hepatocytes were greatly expanded (**Fig. 6A, B**; *n* = 2,055), compared to the control group (*n* = 244). (2) The genes in the WNT-β-catenin pathway were up-regulated in hepatocytes (**Fig. 6C**). These two findings collectively reinforce the hypothesis that, in the liver, *RSPO3* acts in a paracrine manner. Specifically, *RSPO3* is expressed by the endothelial cells to target the adjacent hepatocytes, inducing hepatocyte expansion through the WNT signaling.

**Figure 6.**
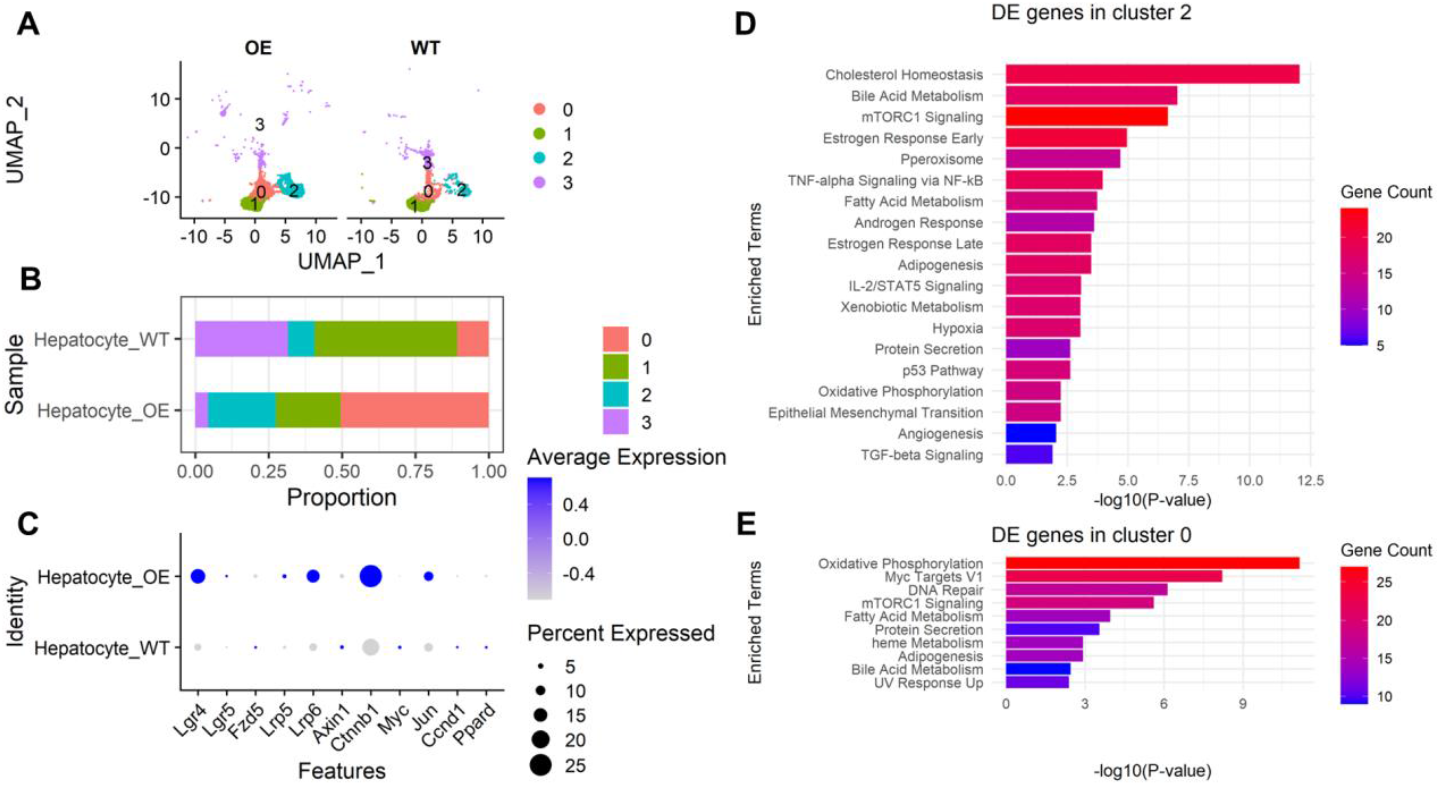
Endothelial factor *Rspo3* can promote MASLD by inducing hepatocyte expansion and affecting fatty acid metabolism via the WNT signaling pathway. (**A**) The hepatocytes are clustered into four groups. (**B**) Compared to hepatocytes of WT, hepatocytes of the ADK-OE group in clusters 0 and 2 are expanded, while the numbers of hepatocytes in clusters 1 and 3 remain relatively stable. (**C**) Genes from the WNT signaling pathway showing enhanced expression in the ADK-OE group. (**D**) Functional enrichment analysis of genes highly expressed in the WT cluster 2. (**E**) Functional enrichment analysis of gene highly expressed in the WT cluster 0.

We extended our investigation to two distinct hepatocyte clusters that showed significant expansion in the ADK-OE group. Our focus was particularly on the WT group to discern the functional differences between these clusters. Using the clusterDE [38], we compared the gene expression profiles of the two hepatocyte clusters under WT conditions (cluster 2 WT vs. cluster 0 WT). Genes highly expressed in the WT cluster 2 were found to be enriched in: (1) fat metabolism-related pathways such as *cholesterol homeostasis, bile acid metabolism*, and *fatty acid metabolism*, (2) the activation of sex hormone-related-pathways such as *estrogen response early*, (3) cancer-related pathways such as *p53 pathway, angiogenesis, mTORC1 signaling, epithelial-mesenchymal transition*, and *TNF-alpha signaling via NF-kB*, and (4) inflammatory response-related pathways such as *IL-2/STAT5 signaling* (**Fig. 6D**). Genes highly expressed in the WT cluster 0 only were enriched fat metabolism-related pathways and cell proliferation pathways such as *adipogenesis* and *Myc targets v1* (**Fig. 6E**). This contrast highlights a key functional distinction between the two clusters. Our results revealed *RSPO3*’s multifaceted roles in liver cell dynamics: promoting hepatocyte expansion, influencing metabolic processes, and liver-specific oncogenic genes through steroid hormone signaling.

## 4. Discussion

In this study, we use a data-driven approach to advance the understanding of how estrogen, *RSPO3*, and body fat distribution interact, and elucidate the implication of this interaction in fatty liver disease and cancer. We highlight three pivotal contributions. First, unveiling the estrogen-*RSPO3* axis. We reveal a crucial link between estrogen and *RSPO3*, which provides novel insights into the sex dimorphism in body fat distribution, and associated cancer disparities. This finding, based on single-cell resolution analysis, establishes a mechanistic connection between hormonal influence and body fat distribution, with its significance in the prevalence of certain cancers. Specifically, we found that estrogen-mediated downregulation of *RSPO3* may inhibit the downstream WNT/β-catenin pathway, which explains the lower susceptibility of females to hepatocellular carcinoma. Second, *RSPO3*’s oncogenic role in the liver. We extend the scope of *RSPO3* from its known association with intestinal cancer to its pivotal role in liver oncogenesis. This novel discovery adds a significant dimension to our understanding of *RSPO3* and opens potential avenues for therapeutic intervention, particularly in the treatment of liver cancer. Third, ADK’s role in fatty liver disease. We demonstrate how *ADK* overexpression contributes to fatty liver disease and obesity by linking it with increased *RSPO3* signaling. This connection not only broadens our understanding of the molecular pathways involved in *ADK*-induced obesity and fatty liver disease but also suggests new targets for treating obesity-related diseases.

## 5. Conclusion

Our study underscores the need for further research. First, deepening understanding of estrogen and *RSPO3* regulation. While we have shown estrogen’s inhibitory effect on *RSPO3*, the underlying mechanisms, particularly the potential epigenetic control involving demethylation, need deeper exploration to fully comprehend these complex interactions. Second, experimental validation of *RSPO3*’s role in the liver. To corroborate our findings on *RSPO3*’s regulatory effect, particularly in ADK-induced obesity, further experiments, such as *RSPO3* knock-down, will be essential to functionally validate and characterize its role in liver conditions. With recent advances in computational methods [39-42] enabling virtual exploration of gene functions, we anticipate that *RSPO3*’s roles will soon be further uncovered.

## 6. Supplementary data

The supplementary data are available within the article.

## Funding

This research was funded by the U.S. Department of Defense (DoD) grant GW200026 for JJC.

## Conflict of interests

The authors declare that they have no competing interests.

## Availability of data and materials

The RNA-seq data from human adipose tissue is from Carithers *et al*. [21] (https://gtexportal.org). Microarray data from the Icelandic Family Adipose (IFA) cohort (*n* = 701) is downloaded from the GEO database (GSE7965). The mouse mammary gland scRNA-seq dataset is downloaded from the GEO database (GSE191219). The 10x Visium spatial transcriptomics data from human colorectal cancer is downloaded from the 10x Genomics website, obtained by Space Ranger 2.0.1 using 10x Genomics Cloud Analysis. The mouse liver NPC scRNA-seq data from Li *et al*. [17] is available from the corresponding author upon reasonable request.

## Ethical Statement

It is not applicable to the scope of this article.

## Authors’ contribution

Q.X. formulated the study and the overall approach. Q.X. performed the analysis with advice from J.J.C and M.C. Q.X., Y.Y, S.G, B.J.C., and G.L. drafted the manuscript.

Corresponding author: Correspondence to Qian Xu.

## Notes

### Competing Interest Statement

The authors have declared no competing interest.

